# Flow modification associated with reduced genetic health of a river-breeding frog, *Rana boylii*

**DOI:** 10.1101/316604

**Authors:** Ryan A. Peek, Sean M. O’Rourke, Michael R. Miller

**Affiliations:** Center for Watershed Sciences, University of California, Davis, CA 95616, USA; Department of Animal Science, University of California, Davis, CA 95616, USA

**Keywords:** Flow alteration, frogs, hydrologic connectivity, hydropower, population genetics, *Rana boylii*, river management, seasonality, Sierra Nevada

## Abstract

River regulation or flow modification—the hydrological alteration of flow by dams and diversions—has been implicated as a cause of fundamental change to downstream aquatic ecosystems. Flow modification changes the patterns and functionality of the natural flow regime, and has the potential to restrict population connectivity and gene flow in river-dependent organisms. Since population connectivity and the maintenance of genetic diversity are fundamental drivers of long-term persistence, understanding the extent flow modification impacts these critical attributes of genetic health is an important goal for long-term conservation. Foothill yellow-legged frogs (*Rana boylii*) were historically abundant throughout many western rivers but have declined since the onset of regulation. However, the extent to which *R. boylii* populations in rivers with altered flow regimes are maintaining connectivity and genetic diversity is unknown. Here we use genetic methods to investigate the impacts of flow alteration on *R. boylii* to explore their potential for long-term persistence under continued flow modification. We found *R. boylii* in rivers with flow modification showed striking patterns of isolation and trajectories of genetic diversity loss relative to unregulated rivers. For example, flow modification explained the greatest amount of variance in population genetic differentiation compared with other covariates including geographic distance. Importantly, patterns of connectivity and genetic diversity loss were observed regardless of flow alteration level but were most prominent in locations with the greatest flow modification intensity. Although our results do not bode well for long-term persistence of *R. boylii* populations under current flow regulation regimes, they do highlight the power of genetic monitoring for assessing population health in aquatic organisms.

## Introduction

River regulation, or the hydrological alteration of flow by dams and diversions, impacts the seasonal and interannual flow variability within a watershed. This flow modification changes the natural flow regime and dramatically alters geomorphic and hydrologic connectivity of watersheds (Poff et al. 2007), which may restrict population connectivity (Schick and Lindley 2007, Shaw et al. 2016) and gene flow (Deiner et al. 2007, Cazé et al. 2016, Thompson et al. 2019). River regulation can change flow frequency, magnitude, duration, timing, and rate of change, which can have significant impacts on aquatic organisms and ecological processes (Poff et al. 2007, Yarnell et al. 2010). Flow modification associated with hydropower generation has been implicated as a cause of fundamental changes to downstream aquatic ecosystems (Power et al. 1996, Bunn and Arthington 2002, Moyle et al. 2011). The hydrological regimes of over half of the world’s largest rivers have been altered by large dams (Nilsson et al. 2005) and only recently has the extent of flow alteration and the associated ecosystem-level impacts been acknowledged (Pringle 2001, Dudgeon et al. 2006, Murchie et al. 2008).

Rivers simultaneously connect and carve the landscapes through which they flow. Rivers provide corridors of connectivity for riparian and aquatic organisms such as fish, amphibians, and macroinvertebrates (Wiens 2002, Pringle 2003), while also acting as physical barriers on the landscape for many terrestrial organisms (Voelker et al. 2013, Cazé et al. 2016). In Mediterranean climates, rivers have strong seasonal patterns associated with cold, wet winters and warm, dry summers. Native aquatic organisms have evolved life histories well adapted to these natural flow patterns, which are both predictable and seasonal (Yarnell et al. 2010, Tonkin et al. 2017). Thus, changes to temporal and spatial abiotic river processes caused by river regulation can have a substantial impact on biotic communities. The negative effects of flow alteration on migration and loss of spawning habitat (Lind et al. 1996, Fuller et al. 2011, Kupferberg et al. 2012, Rolls and Bond 2017), reductions in population abundances and diversity (Lind et al. 1996, Zhong and Power 1996, Vörösmarty et al. 2010, Fuller et al. 2011, Scribner et al. 2016, Sabo et al. 2017, Guzy et al. 2018), and fragmentation (Zhong and Power 1996, Vörösmarty et al. 2010, Werth et al. 2014, Scribner et al. 2016, Sabo et al. 2017, Guzy et al. 2018) have been well documented. However, most rivers have not been regulated for long periods (e.g., typically less than 100 years) compared to the time aquatic organisms had to adapt to pre-anthropogenic flow conditions. In flow-altered rivers that organisms still occupy, it remains unknown whether populations can persist long-term with continued flow modification. In other words, while some species may have persisted since flow regulation began in a system (e.g., typically four to five decades), this does not necessarily mean these populations will persist into the future under current altered flow regimes. Thus, exploring the potential for long-term persistence of populations under different altered flow regimes is a crucial component for guiding conservation efforts, yet it remains a significant gap.

One tool that can help address this gap is the integration of genetics and hydrology to better assess the impact of flow alteration on aquatic organisms (Scribner et al. 2016). Although aquatic organisms are often difficult to count and monitor by conventional methods, genetic monitoring can be a powerful tool to assess population health by revealing factors such as genetic fragmentation and population declines. It is widely recognized that reductions in population connectivity can increase inbreeding, leading to a potential “extinction vortex” (Gilpin and Soule 1986), yet there is limited understanding of how flow alteration may impair the processes crucial for maintenance of genetic variation and thus adaptive capacity. In addition, there is a pressing need for more effective and flexible watershed management tools, particularly in relation to monitoring aquatic populations, and assessment of environmental flows (Grantham et al. 2010, Yarnell et al. 2020). Thus, population genetics could be a powerful tool to understand the influence of different flow regimes on population health and this information could facilitate improved flow management to better protect aquatic populations.

The river-breeding foothill yellow-legged frog (*Rana boylii*) historically occurred in lower and mid-elevation streams and rivers from Southern Oregon to northern Baja California west of the Sierra-Cascade crest (Stebbins 2003). *Rana boylii* are intimately linked with river hydrology because they have evolved to spawn in synchrony with natural flow cues associated with seasonal spring snowmelt or rain recession periods (Kupferberg 1996, Yarnell et al. 2010, 2016, Bondi et al. 2013). However, population declines have been documented across the former range of this species, particularly in southern California and the Sierra Nevada where it has been extirpated from approximately 50 percent of its historical range (Jennings and Hayes 1994, Davidson et al. 2002). In California, particularly in the Sierra Nevada, flow alteration may be a significant environmental stressor (Lind et al. 1996, Kupferberg et al. 2012). Regulated river reaches typically alter flows by augmenting or diverting winter and spring runoff, thereby disrupting natural flow regimes and reducing or eliminating flow cues.

Aseasonal flow fluctuation from flow alteration can scour (i.e., detach from substrate) or desiccate *R. boylii* egg masses, and the loss of clutches may have a significant demographic impact because only one egg mass is laid per year. In many regulated rivers in the Sierra Nevada, *R. boylii* populations are now restricted to small unregulated tributaries flowing into the regulated mainstem (Peek 2018).

Here, we investigate the impacts of flow modification on genetic health of *R. boylii* populations across three different flow regimes. Given that population connectivity and genetic diversity are known to be play critical roles in long-term species persistence, our primary questions were:

1. How has flow alteration impacted seasonality and predictability of flow patterns in regulated river reaches?
2. Do populations of *R. boylii* in flow altered river reaches have less genetic diversity and decreased measures of genetic connectivity than populations in unregulated river reaches?
3. Are metrics of connectivity and diversity in *R. boylii* associated with different levels of flow alteration?

Our goal was to assess the genetic health of *R. boylii* under different flow regimes to better inform the potential for long-term persistence with flow alteration. Addressing this issue will help to inform management and conservation efforts for *R. boylii*, through evaluation and prioritization of conservation actions for populations with decreased genetic health. In addition, our approach demonstrates the potential utility of genetics for future conservation monitoring and assessment efforts in aquatic species in altered systems.

## Material And Methods

To begin investigating the impact of river regulation on *R. boylii*, we collected frog tissue and buccal samples from 30 locations in six rivers representing three different flow alteration levels associated with hydropower generation (Table 1). The three flow regimes assessed were: 1) hydropeaking, where flows are pulsed on most days from late spring through fall to provide electricity during peak-use hours and for recreational whitewater rafting; 2) bypass, which diverts river flows from an upstream portion of the basin to the downstream power generation facilities; and 3) unregulated, a largely natural flow regime where no upstream controls exist to regulate flows (Figure 1A). Flow data were obtained for each river reach using proximal USGS gaging stations (Data S1). We analyzed *R. boylii* samples from sites in three major watersheds (Yuba, Bear, and American) in the northern Sierra Nevada of California (Figure 1A; Table 1, Data S2). The six study rivers share a similar Mediterranean climate, underlying geology, watershed aspect (west-slope), stream morphology (riffle-pool), and vegetative communities, but differ in the intensity of flow alteration (Steel et al. 2017). Although flow alteration occurs in all three of the study watersheds, both the North Yuba and North Fork (NF) American are unregulated whereas the Middle Fork (MF) American is the only river that has a hydropeaking flow regime (Figure 1B, Figure 1C).

**Table 1.** Locality information for population genomic analysis of *R. boylii* in the Yuba, Bear, and American Watersheds.

| Locality | Site ID | River | Watershed | Regulation Type | Lat. (WGS84) | Long. (WGS84) | Elev. (m) | NHD<sup>+</sup> Stream Order | NHD Total Drainage Area (sq. km) |
| --- | --- | --- | --- | --- | --- | --- | --- | --- | --- |
| American Canyon | 1 | MF American | American | Hydropeaking | 38.93396 | -120.94356 | 240 | 2 | 9 |
| Gas Canyon | 2 | MF American | American | Hydropeaking | 38.96651 | -120.93252 | 241 | 1 | 13 |
| Todd Creek | 3 | MF American | American | Hydropeaking | 38.96385 | -120.92158 | 367 | 2 | 10 |
| NFMFA-Skunk Canyon | 4 | MF American | American | Hydropeaking | 39.02237 | -120.73693 | 521 | 2 | 6 |
| Rubicon-US Powerhouse | 5 | MF American | American | Bypass | 38.99928 | -120.72333 | 360 | 5 | 816 |
| Rubicon-Long Canyon | 6 | MF American | American | Bypass | 38.98887 | -120.68996 | 415 | 5 | 806 |
| Ponderosa Bridge | 7 | NF American | American | Unregulated | 38.99994 | -120.9406 | 240 | 5 | 857 |
| Bunch Canyon | 8 | NF American | American | Unregulated | 39.03762 | -120.91028 | 286 | 3 | 27 |
| Shirrtail Creek | 9 | NF American | American | Unregulated | 39.04446 | -120.89937 | 525 | 4 | 141 |
| Indian Creek | 10 | NF American | American | Unregulated | 39.05665 | -120.90854 | 296 | 2 | 24 |
| Slaughter Ravine | 11 | NF American | American | Unregulated | 39.09865 | -120.92548 | 356 | 2 | 6 |
| Robbers Ravine | 12 | NF American | American | Unregulated | 39.10451 | -120.92672 | 400 | 1 | 4 |
| Iowa Hill Mainstem | 13 | NF American | American | Unregulated | 39.11115 | -120.91684 | 386 | 4 | 605 |
| Euchre Bar | 14 | NF American | American | Unregulated | 39.18492 | -120.76204 | 579 | 4 | 508 |
| Sailor Canyon | 15 | NF American | American | Unregulated | 39.21694 | -120.49595 | 1005 | 3 | 166 |
| Greenhorn Creek | 16 | Bear | Bear | Bypass | 39.23206 | -120.90189 | 820 | 2 | 11 |
| Steep Hollow Creek DS | 17 | Bear | Bear | Bypass | 39.20231 | -120.87537 | 736 | 2 | 52 |
| Hawkins Ravine | 18 | Bear | Bear | Bypass | 39.18833 | -120.8981 | 706 | 2 | 4 |
| Steep Hollow Creek US | 19 | Bear | Bear | Bypass | 39.19444 | -120.88778 | 704 | 2 | 45 |
| Chicago Powerhouse | 20 | Bear | Bear | Bypass | 39.17484 | -120.89978 | 665 | 4 | 136 |
| Shady Creek | 21 | South Yuba | Yuba | Bypass | 39.35433 | -121.059 | 675 | 2 | 15 |
| Spring Creek | 22 | South Yuba | Yuba | Bypass | 39.33233 | -120.989 | 595 | 3 | 24 |
| Rock Creek | 23 | South Yuba | Yuba | Bypass | 39.32983 | -120.98634 | 593 | 4 | 710 |
| Missouri Canyon | 24 | South Yuba | Yuba | Bypass | 39.36096 | -120.88138 | 1094 | 2 | 5 |
| Logan Creek | 25 | South Yuba | Yuba | Bypass | 39.36914 | -120.85258 | 1201 | 1 | 5 |
| US Canyon Creek | 26 | South Yuba | Yuba | Bypass | 39.35387 | -120.73419 | 889 | 4 | 365 |
| Oregon Creek | 27 | Middle Yuba | Yuba | Bypass | 39.44188 | -121.05753 | 620 | 4 | 375 |

| **Locality** | **Site ID** | **River** | **Watershed** | **Regulation Type** | **Lat. (WGS84)** | **Long. (WGS84)** | **Elev. (m)** | **NHD<sup>†</sup> Stream Order** | **NHD Total Drainage Area (sq. km)** |
| --- | --- | --- | --- | --- | --- | --- | --- | --- | --- |
| US Our House Dam | 28 | Middle Yuba | Yuba | Bypass | 39.41305 | -120.99035 | 624 | 4 | 375 |
| Rocky Rest Mainstem | 29 | North Yuba | Yuba | Unregulated | 39.51191 | -120.97738 | 704 | 5 | 669 |
| Slate Creek | 30 | North Yuba | Yuba | Bypass | 39.689128 | -120.93889 | 1330 | 3 | 58.7 |

<sup>†</sup>NHD refers to the National Hydrography Dataset by USGS (U.S. Geological Survey, National Hydrography
Dataset, Digital data, accessed, August 2017).

**Figure 1.**
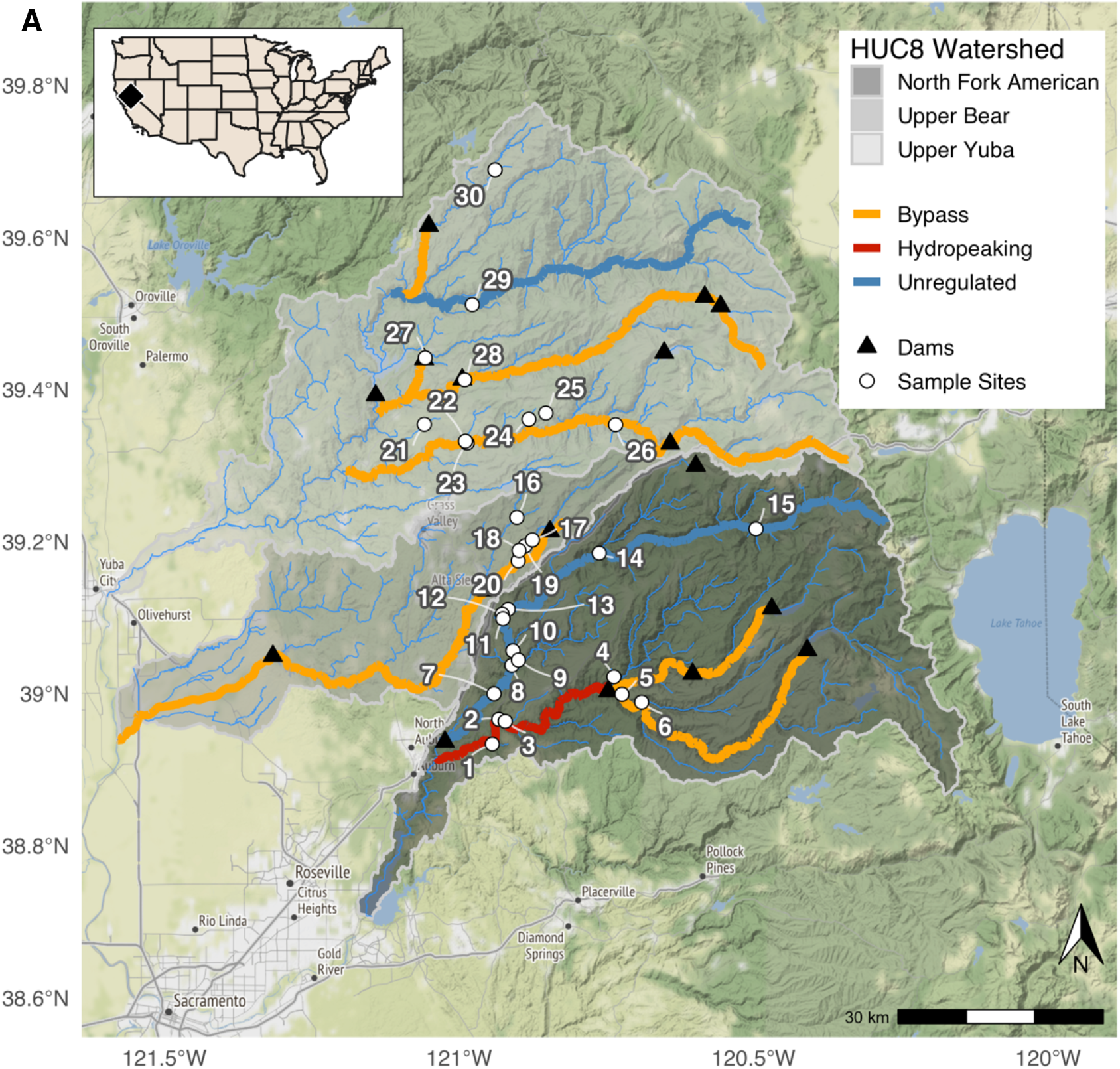
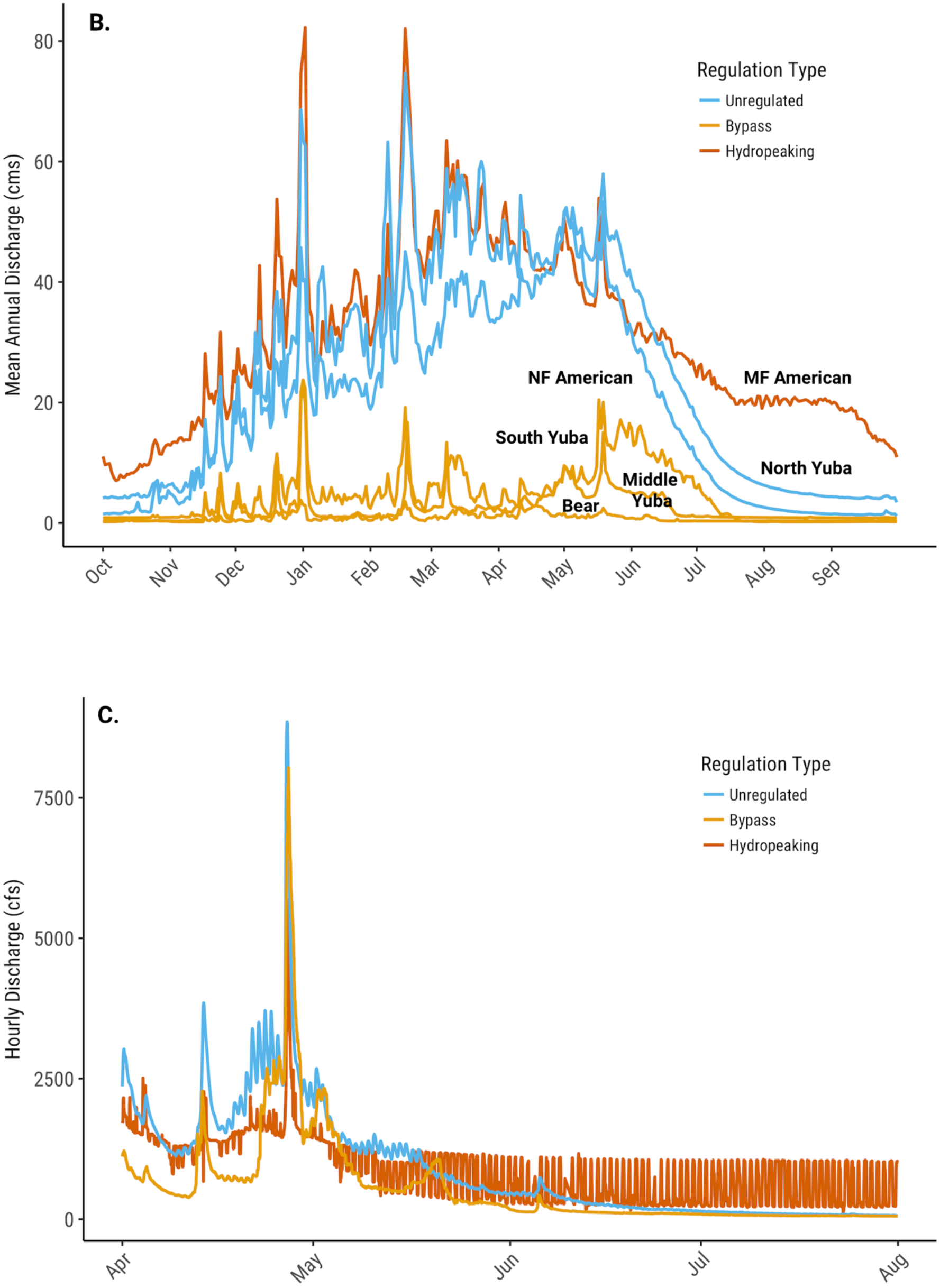
Sampling locations and flow characteristics. A) Map of sampling locations across three watersheds. B) Comparison of annual mean daily discharge from 1981–2016 for three flow types. C) Comparison of hourly discharge in three different flow regimes in April through July 2012, Bypass (South Yuba), Hydropeaking (Middle Fork American), and Unregulated (North Fork American). See Table S1 for USGS gaging station information.

### Assessment of hydrologic impacts of river regulation on patterns of seasonality

To evaluate how flow alteration has impacted seasonal flow patterns in regulated river reaches, we used a wavelet analysis to derive metrics of predictability and Colwell’s information-theoretic metrics to define seasonality following methods outlined in Tonkin et al. (2017). Seasonality—how much the environment varies during the course of a single year—was measured using Colwell’s M/P metric, which is the contingency (M) divided by within season predictability (P) (Colwell 1974). Predictability was measured using the standardized Morlet wavelet power at the 12-month time interval across the entire data series (Torrence and Compo 1998). Daily average flow data from all available gages within the study area were used to compare the inter-annual (wavelet) and intra-annual (Colwell’s indices) flow patterns in each of the study reaches (Data S1, Data S3). In regulated reaches where data was available, metrics were estimated from the full period of record prior to initiation of flow alteration (pre-regulation) as well as from current altered flow regimes.

### Sample collection and DNA extraction

To evaluate genetic diversity and connectivity in *R. boylii* populations, a total of 345 tadpole tail clips, buccal swabs, and tissue samples were compiled. The samples were collected between 1992 and 2017, from watersheds across the west slope of the Sierra Nevada (Data S2). Field sampling was conducted following Heyer et al. (1994), under CDFW SCP Permit #0006881, with IACUC protocol #19327. Individual post-metamorphic frogs were buccal-swabbed following established protocols (Goldberg et al. 2003, Pidancier et al. 2003, Broquet et al. 2007). Each post-metamorphic individual was comprehensively swabbed underneath tongue and inside of both cheeks for approximately 30 sec to one minute. Swabs were air dried for approximately five minutes and placed in 1.5 mL microcentrifuge tubes while in the field or placed in lysis buffer (Goldberg et al., 2003). Dried samples were stored in the laboratory at −80°C until DNA extraction. Where possible, tail clips from tadpole larvae were collected, and tadpoles greater than 15 mm total length were targeted (Parris et al., 2010; Wilbur & Semlitsch, 1990). One clip was taken per individual tadpole and dried on Whatman filter paper (grade 1) or placed in 95% ethanol and stored at room temperature. DNA was extracted from samples in ethanol and lysis buffer using Qiagen DNeasy Blood & Tissue kits following manufacturer protocol and stored at −20°C. DNA was extracted from dried buccal swabs and tail clips using an Ampure magnetic bead-based protocol (Ali et al., 2016) and stored at −20°C.

### RAD Sequencing and RAD-Capture (Rapture) bait design

To produce a genomic resource for frog species with large genome sizes, we interrogated a significant fraction of the *R. boylii* genome using RAD sequencing with SbfI (Miller et al. 2007, Baird et al. 2008, Ali et al. 2016). Paired-end Illumina sequence data were generated using 24 *R. boylii* individuals (Data S4) and de novo locus discovery and contig extension were carried out as previously described (Miller et al. 2012, Sağlam et al. 2016, Peek et al. 2019) using the alignment program Novoalign and the PRICE assembler (Ruby et al. 2013). This resulted in a set of 77,544 RAD contigs ranging from 300 to 800 bp which served as a de novo partial reference assembly for all subsequent downstream analyses (Data S5). We next removed loci with five or more SNPs to reduce potential paralogs and chimeras, and randomly selected 10,000 loci from the remaining subset. Of these 10,000 loci, 8,533 were successfully designed into 120 bp RAD capture baits by Arbor Biosciences (Data S6).

### Rapture Sequencing and Genomic Analysis

A total of 345 individual frog samples were prepared for sequencing following the RAD Capture (Rapture) methods outlined in Ali et al. (2016) and detailed in Peek et al. (2019). We generated RAD libraries from the samples, quantified the libraries using a Fragment Analyzer (Agilent Technologies, Santa Clara, CA) and pooled them, performed capture on the pooled library with the 120 bp baits described above, and sequenced the resulting Rapture library with paired-end Illumina sequencing.

The mean number of filtered alignments across all 345 samples was 324,928, and sequence coverage depth was measured at the mid-point (i.e., base-pair 60) of each 120 bp Rapture bait locus using the depth function in SAMtools (Li et al. 2009, Li 2011). For downstream analysis, we selected individuals that had greater than 100,000 alignments (n=277), which provided sufficient data to investigate population genetic attributes at broad and fine geographic scales (Table 2, Data S2). *Rana boylii* are cryptic, and often occur in low densities within the study area. Thus, we retained a minimum of three individuals per site, and the mean number of samples per site was approximately nine (Table 2). With genomic data, population genetic parameters can be accurately estimated from even low sample numbers (Hotaling et al. 2018), and genomic analyses in non-model organism often use fewer loci than analyzed here (Narum et al. 2013). Thus, the sequence data we obtained should be appropriate for population genetic analyses across our study area.

**Table 2.** Sampling information for population genomic analysis of *R. boylii*, the number of individuals (n) is given for the total number sequenced per location and the number of individuals that were retained after filtering (see methods) across the 8,533 baits.

| Locality | Site ID | n initial | n retained |
| --- | --- | --- | --- |
| American Canyon | 1 | 16 | 6 |
| Gas Canyon | 2 | 6 | 6 |
| Todd Creek | 3 | 11 | 9 |
| NFMFA-Skunk Canyon | 4 | 18 | 18 |
| Rubicon-US Powerhouse | 5 | 11 | 11 |
| Rubicon-Long Canyon | 6 | 9 | 8 |
| Ponderosa Bridge | 7 | 5 | 5 |
| Bunch Canyon | 8 | 15 | 14 |
| Shirttail Creek | 9 | 16 | 15 |
| Indian Creek | 10 | 12 | 11 |
| Slaughter Ravine | 11 | 8 | 8 |
| Robbers Ravine | 12 | 30 | 11 |
| Iowa Hill Mainstem | 13 | 36 | 30 |
| Euchre Bar | 14 | 13 | 11 |
| Sailor Canyon | 15 | 8 | 5 |
| Greenhorn Creek | 16 | 15 | 6 |
| Steep Hollow Creek DS | 17 | 13 | 12 |
| Hawkins Ravine | 18 | 3 | 3 |
| Steep Hollow Creek US | 19 | 7 | 6 |
| Chicago Powerhouse | 20 | 6 | 6 |
| Shady Creek | 21 | 14 | 12 |
| Spring Creek | 22 | 4 | 4 |
| Rock Creek | 23 | 3 | 3 |
| Missouri Canyon | 24 | 8 | 6 |
| Logan Creek | 25 | 5 | 4 |
| US Canyon Creek | 26 | 6 | 6 |
| Oregon Creek | 27 | 15 | 13 |
| US Our House Dam | 28 | 13 | 12 |
| Rocky Rest Mainstem | 29 | 15 | 12 |
| Slate Creek | 30 | 4 | 4 |

The probabilistic framework implemented in Analysis of Next Generation Sequencing Data (ANGSD) (Korneliussen et al. 2014) was used for all population genetic analyses as it does not require calling genotypes and is suitable for low-coverage sequencing data (Korneliussen et al. 2013, Fumagalli et al. 2013). ANGSD analyses were conducted following details from Peek et al. (2019).

### Principal component analysis to assess population structure and connectivity

To assess *R. boylii* population structure across the collection locations, we used ANGSD (Korneliussen et al. 2014) to perform principal component analysis (PCA). Settings used in ANGSD for PCA to identify polymorphic sites for PCA included a SNP_pval of 1×10^−6^, inferring major and minor alleles (doMajorMinor 1), estimating allele frequencies (doMaf 2) (Kim et al. 2011), retaining SNPs with a minor allele frequency of at least 0.05 (minMaf), genotype posterior probabilities were calculated with a uniform prior (doPost 2), and the doIBS 1 and doCov 1 options were used to generate PCA data. Principal components (PC) summarizing population structure were derived from classic eigenvalue decomposition and were visualized using the {ggplot2} package (Wickham 2016) in R (R Core Team 2017).

### Genetic differentiation and diversity estimates

To investigate patterns of genetic diversity and the overall trajectory (stable, increasing, or decreasing) of diversity in a population, we summarized patterns of genetic variation using two estimators of θ (4Nμ): Tajima’s θ (θ_π_) is based on the average number of pairwise differences (Tajima 1983), and Watterson’s θ (θ_S_) is based on the number of segregating sites (Watterson 1975). Estimates of θ were made using the empirical Bayes method in ANGSD (Korneliussen et al. 2014) with the site frequency spectrum (SFS) to calculate each statistic for each site, which were then averaged to obtain a single value for each statistic (Korneliussen et al. 2013). These estimators are influenced by the demographic history of a population and provide information on the trajectory of changes in genetic diversity (Tajima 1989, Wakeley 2009). When genetic diversity has been stable, these estimates should be equal; when genetic diversity has been decreasing, θ_π_ > θ_S_ and when genetic diversity has been increasing, θ_π_ < θ_S_. Scaled Δθ ([θ_π_ − θ_S_] / ([θ_π_ + θ_S_] / 2)) was used to assess trajectories of genetic diversity in each population.

To assess how patterns of genetic differentiation and connectivity are associated with flow alteration in the study, we calculated pairwise F_ST_ (Wright 1943) between all collection locations within a river for all six rivers. Genome-wide F_ST_ between location pairs was estimated by first calculating a site frequency spectrum (SFS) for each population (doSaf) (Nielsen et al. 2012) with ANGSD. The two-dimensional SFS and global F_ST_ between each population pair were then estimated using realSFS (Korneliussen et al. 2014). We then plotted the scaled pairwise F_ST_ (F_ST_ / [1-F_ST_]) (Rousset 1997) for each location against the river distance (the distance along the river network from each collection location to every other location within that study river). We calculated the river distances (distance along river network) between locations within watersheds using the {riverdist} package in R (Tyers 2017). These values were plotted and a generalized linear model was fitted (Scaled F_ST_ ~ River Distance) in R (R Core Team 2017). Furthermore, each location was assigned flow alteration type based on either the regulation type at the mainstem sample location or at the nearest mainstem; for pairwise comparisons the type was assigned based on whichever was greatest (i.e., hydropeaking is of greater intensity than bypass).

### Boosted regression tree modeling of variance in F_ST_

We used boosted regression tree (BRT) models with the R packages {gbm} (Ridgeway 2015) and {dismo} (Hijmans et al. 2017) to assess the relative influence of flow alteration type as compared to other environmental covariates. Boosted regression trees (BRT) are suitable frameworks for large and complex ecological datasets because they do not assume normality, nor linear relationships between predictor and response variables and they ignore non-informative predictor variables (Graham et al. 2008, Steel et al. 2017). BRTs use iterative boosting algorithms to combine simple decision trees to improve model performance (De’ath 2007) and provide a robust alternative to many traditional statistical methods (Phillips et al. 2006, Guisan et al. 2007). BRTs assess the relative impact of modeled variables by calculating the number of times a variable is selected for splitting a tree across all folds of the cross validation. Following Steel *et al.* 2017, estimates of relative influence for each predictor variable were used to evaluate the relative contribution a variable had in predicting the response. To evaluate the relative influence of covariates on F_ST_, models were trained using river distance (km), elevation (m), upstream drainage area (km^2^), Strahler stream order, and number of samples per location. Stream segment data on elevation, length, slope, stream order, and drainage area were derived from NHD Plus attributes (U.S. Geological Survey, National Hydrography Dataset, Digital data, accessed, August 2017 at http://nhd.usgs.gov/data.html). In addition, scaled Δθ was included to assess the effect of genetic diversity trajectory on F_ST_ across regulation types.

Model training and fitting were conducted following methods previously described in Steel et al. (2017). To reduce overfitting, the learning rate (also known as the shrinking rate) was set to 0.001. Stochastic gradient boosting was utilized to reduce prediction error (De’ath 2007) and the fraction of training data sampled to build each tree was 0.75, within the range as recommended by (Brown et al. 2012). Tree complexity was set to three to allow for second and third order interaction effects. The minimum number of observations required in the final nodes of each tree was three. A ten-fold cross-validation technique allowed us to determine the number of trees at which prediction error was minimized using the cross-validation deviance. Model performance was evaluated using the minimum estimated cross-validation deviance which maximized the estimated deviance explained.

## Results

### Flow alteration reduces the strength and predictability of seasonality patterns

Flow patterns between flow-altered and unregulated river reaches could be clearly differentiated by flow regime (hydropeaking, bypass, unregulated). Wavelet analysis and Colwell’s M/P metric indicated flow alteration reduced seasonality and predictability of flow patterns in regulated (hydropeaking and bypass) river reaches (Figure 1B, 1C, 2). These patterns were distinctly different before and after flow alteration began in regulated study reaches (Figure 2, Data S3). In all flow-altered reaches, seasonality was reduced compared to pre-regulation flow patterns, and predictability of these flow patterns was also reduced in flow-altered data (Figure 2).

**Figure 2.**
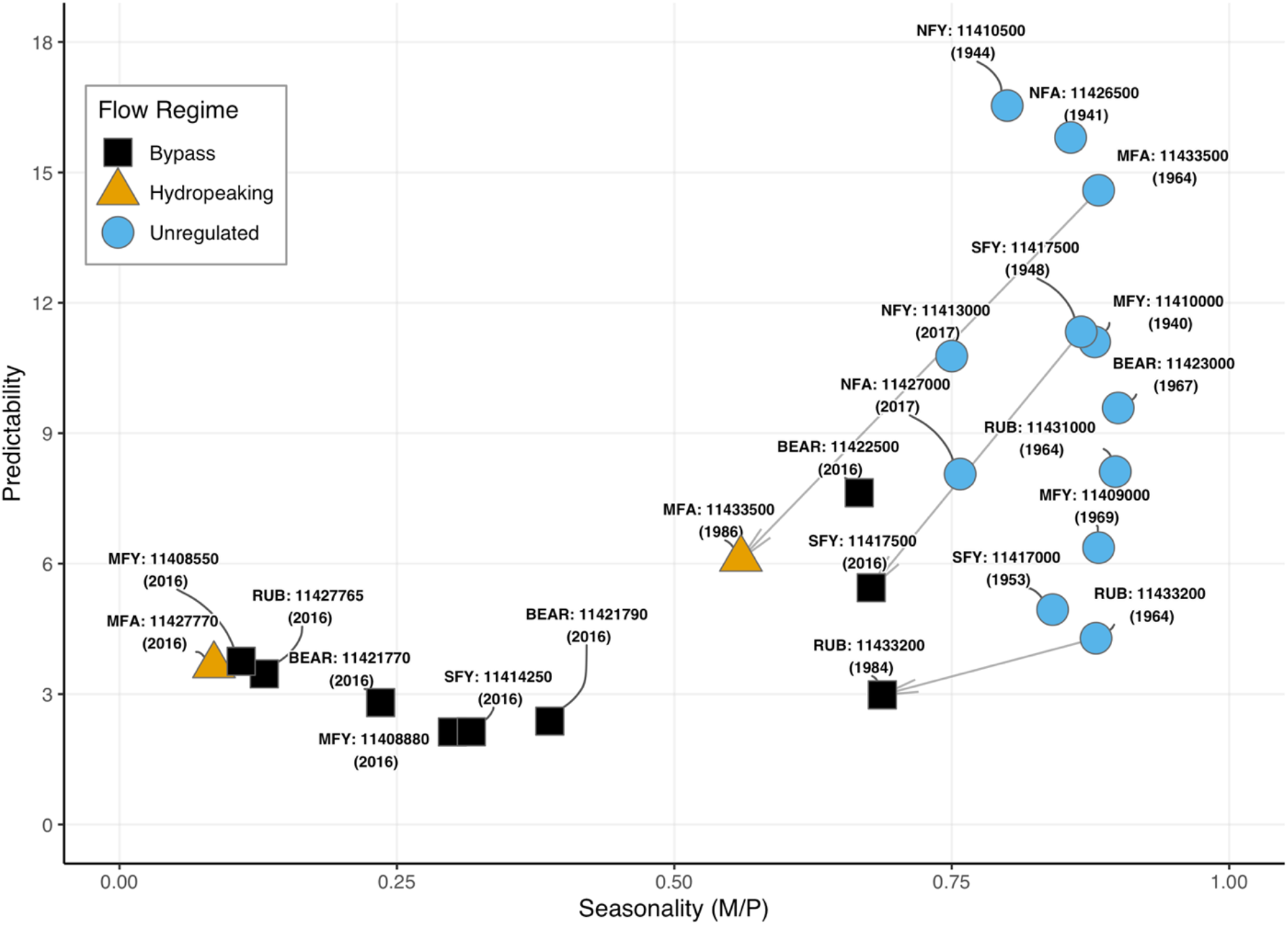
A biplot of seasonality (Colwell’s M/P [contingency/predictability] versus predictability (average wavelet power at the 12-month time period following Tonkin et al. (2017). Each point is labeled with the river acronym and USGS gage ID, followed by the end year of the flow dataset used (See Table S3). Data from the same gage but from two time periods (before flow alteration and with contemporary alteration) are connected by arrows.

### Anomalous genetic pattern in highly regulated reach of Middle Fork American watershed

To assess *R. boylii* population structure across the collection locations, we used principal component analysis to provide a dimensionless comparison of all samples. The first two principal components revealed four main groups corresponding to the Yuba, Bear, North Fork (NF) American, and Middle Fork (MF) American samples (Figure 3A). Unlike the Yuba watershed where all rivers clustered as one group, the two rivers within the American watershed (the NF American and MF American) were separated by both PC1 and PC2. Although the NF American watershed clustered closely with the adjacent Bear watershed, the MF American showed a surprisingly high degree of genetic differentiation from other locations (Figure 3A). These data suggest that there is less genetic differentiation between the NF American and the Bear watersheds, than between the NF and MF American watersheds. We conclude that measurements of overall genetic differentiation in *R. boylii* from our study area largely conform to watershed and geographic expectations, with the exception of the American watershed, which shows a high degree of genetic differentiation between the North (unregulated) and Middle (hydropeaking) Forks.

**Figure 3.**
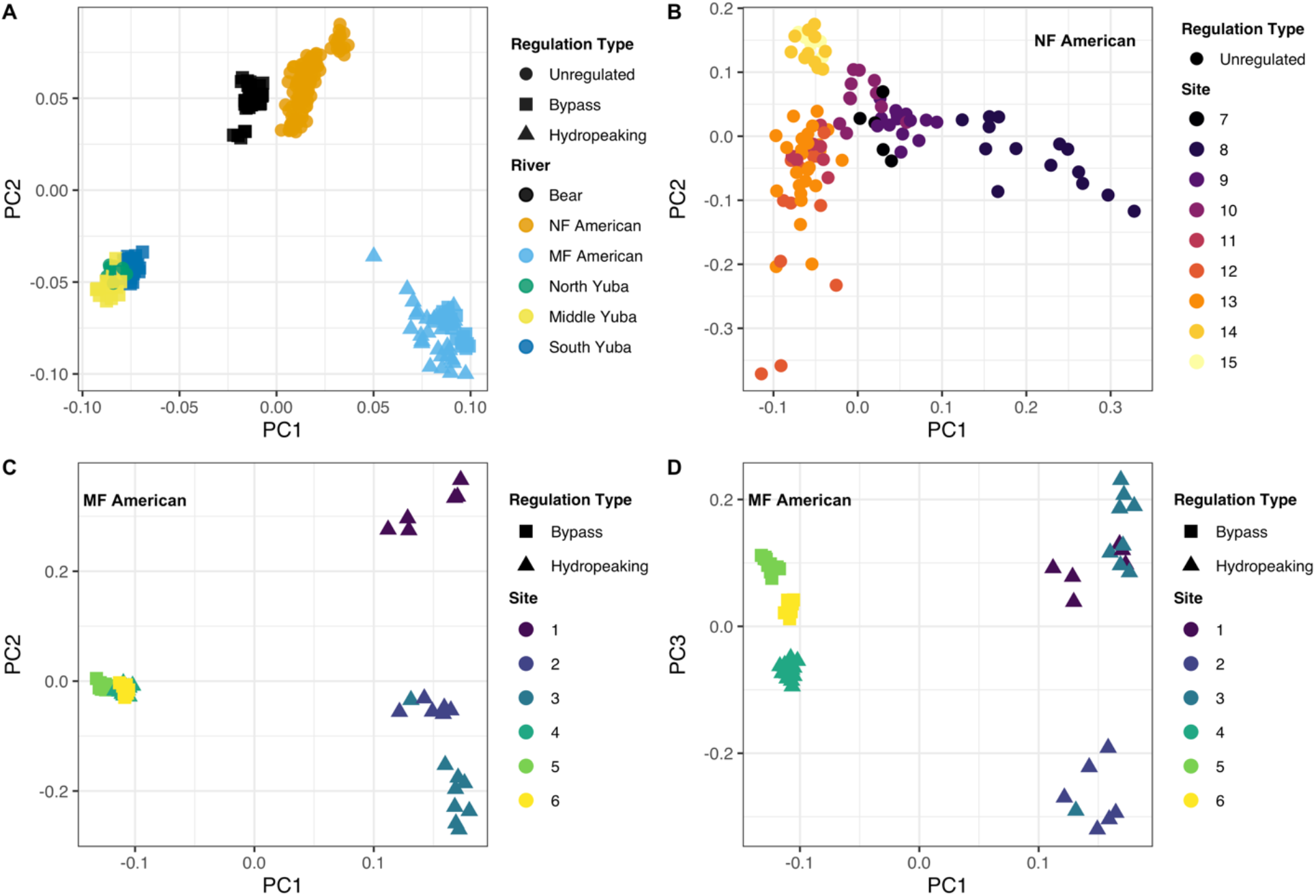
Principal component analysis of Rapture sequencing data. A) Northern Sierra Nevada watersheds; B) Unregulated NF American; C) and D) Hydropeaking MF American Reach.

To further investigate patterns of genetic variation within the American Watershed, we performed two PCAs, one on samples from the NF American, and the other on samples from the MF American. The PCA of the NF American showed minimal differentiation among locations, with different study sites blending together and weak patterns of population structure (Figure 3B). In contrast, PCA of the MF American showed strong differentiation between sites (Figures 3C, 3D). The MF American PCA completely resolved all sites, with the first component (PC1) strongly differentiating the samples in the hydropeaking reach from all other sites in the MF American. This pattern may be due to the differential river regulation between the two rivers; the NF American is unregulated and has weak PCA differentiation, whereas the MF American has a higher level of river regulation and all sites form distinct genetic clusters, indicative of reduced gene flow among sites within the MF American.

### Flow regime is the strongest predictor of genetic isolation with R. boylii in the Northern Sierra

To explore the relationship between the flow alteration type and genetic differentiation across the entire study area, we plotted scaled F_ST_ against river distance between sampling sites for each flow alteration type assessed. While there was a clear relationship between scaled F_ST_ and river distance (as shown by the slope of regression lines in Figure 4A), there was a striking pattern of elevated F_ST_ by flow alteration type (Figure 4A, Data S7). Even the bypass flow alteration type showed a distinct pattern of elevated F_ST_. For instance, sites in hydropeaking flow regimes separated by less than 10km had scaled F_ST_ values comparable to unregulated locations separated by river distances over 50 km. Hydropeaking was the most extreme pattern of the three flow alteration types and showed highly elevated F_ST_ values compared with the unregulated and bypass flow types. The centroid F_ST_ of each flow type increased with increasing flow alteration intensity from unregulated (centroid F_ST_=0.144, se=0.007) to bypass (centroid F_ST_=0.218, se=0.012) to hydropeaking (centroid F_ST_=0.284, se=0.017). The centroid F_ST_ for hydropeaking was nearly double compared with unregulated centroid F_ST_. Furthermore, the centroid of the river distance between site pairs was comparable between flow alteration types (unregulated=20.9 km, bypass=21.4 km, hydropeaking=21.6 km), further supporting patterns of genetic differentiation are associated with flow alteration and not solely isolation by distance. This suggests a greater degree of isolation within sites in reaches with altered flow regimes compared with *R. boylii* populations in unregulated reaches, as larger F_ST_ values represent reductions in heterozygosity due to population subdivision (Slatkin 1987). We conclude *R. boylii* in rivers with altered flow regimes show patterns of greater population isolation and elevated F_ST_, (i.e., reduced heterozygosity) compared to populations in unregulated locations.

**Figure 4.**
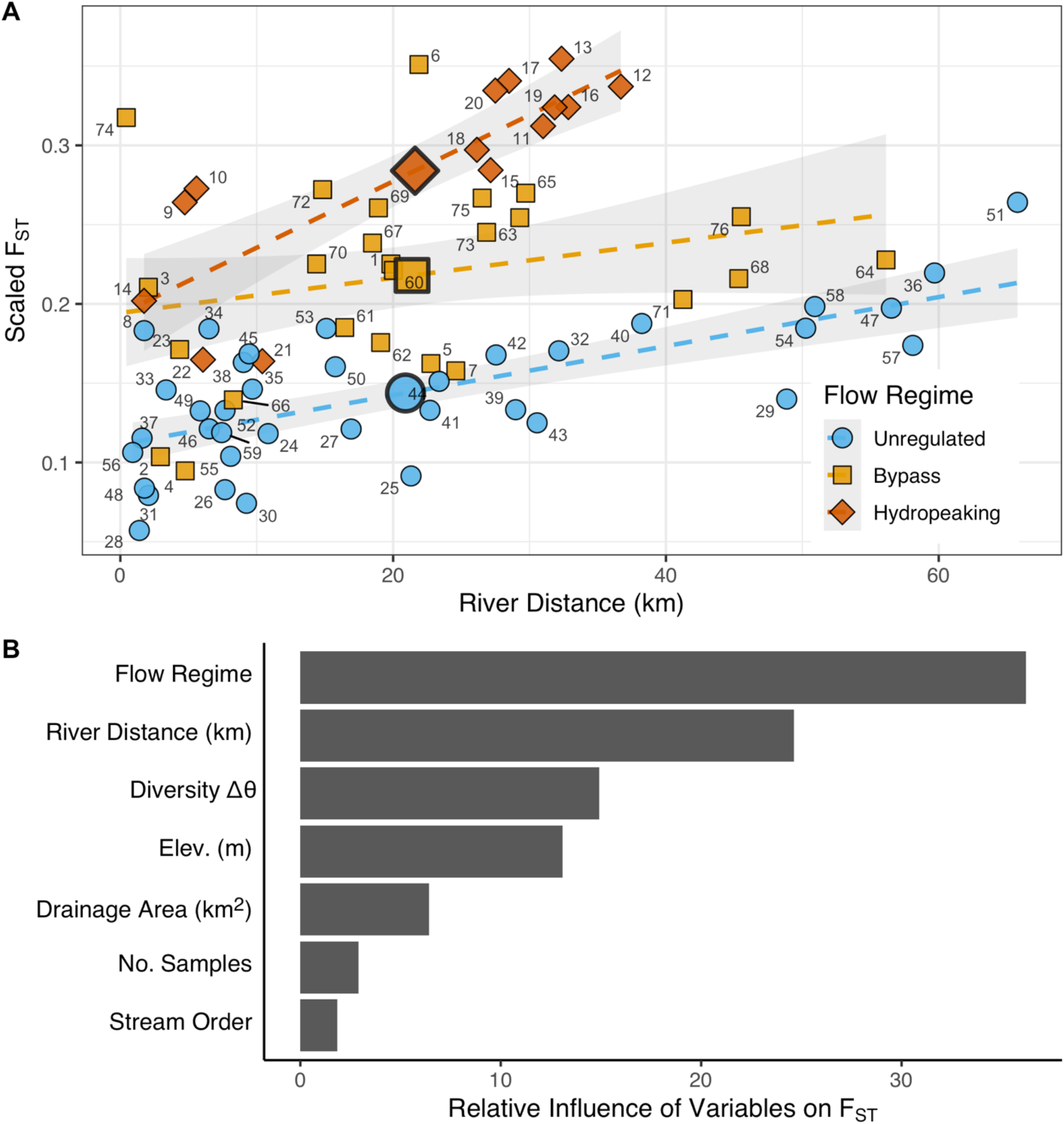
Relationship between river distance and genetic differentiation in *R. boylii*. A) pairwise scaled F_ST_ vs. river distance (km) for each location pair, centroid of each flow alteration type delineated as large point with black outline. Numbers on points correspond to “sitepairID” in Data S7; B) Relative influence of covariates on scaled F_ST_ from boosted regression tree models.

To investigate the relative influence of flow alteration compared to other covariates such as river distance on genetic differentiation (i.e. F_ST_), we used boosted regression tree (BRT) modeling. Covariates included flow regime alteration type, river distance, watershed variables derived from National hydrology data (NHD), topographic data, and genetic diversity trajectory (i.e., scaled Δθ). We found flow alteration explained the greatest amount of variance in scaled F_ST_ (Figure 4B). Thus, flow alteration has a larger relative influence on genetic differentiation than river distance between sampling locations in altered flow reaches. We conclude there is a pattern of elevated F_ST_ between populations in reaches with altered flow.

### Flow alteration is strongly correlated with decreasing genetic diversity in R. boylii

To investigate the association between flow and genetic diversity trajectory(whether genetic diversity is stable, increasing, or decreasing), we summarized patterns of genetic variation using two estimators of θ (4Nμ): Tajima’s θ (θ_π_) and Watterson’s θ (θ_S_). We found zero populations sampled within flow altered watersheds had evidence of increasing genetic diversity (i.e., a θ_π_ − θ_S_ and scaled Δθ ([θ_π_ − θ_S_] / mean θ)) that was less than zero) (Figure 5A). The altered flow locations showed a clear trajectory of genetic diversity loss (Figure 5A, 5B). Three of the four hydropeaking locations had the highest values of scaled Δθ, and the global mean was significantly different from other flow alteration types. Although some tributary populations within unregulated watersheds also showed signs of genetic diversity loss, the mean genetic diversity trajectory at unregulated locations was largely stable (Figure 5B). This indicates many populations in the northern Sierra Nevada which are already limited in number, are losing genetic variation, and flow alteration is exacerbating these patterns. We conclude there is evidence of recent genetic diversity loss across populations in the flow altered river systems, regardless of alteration type.

**Figure 5.**
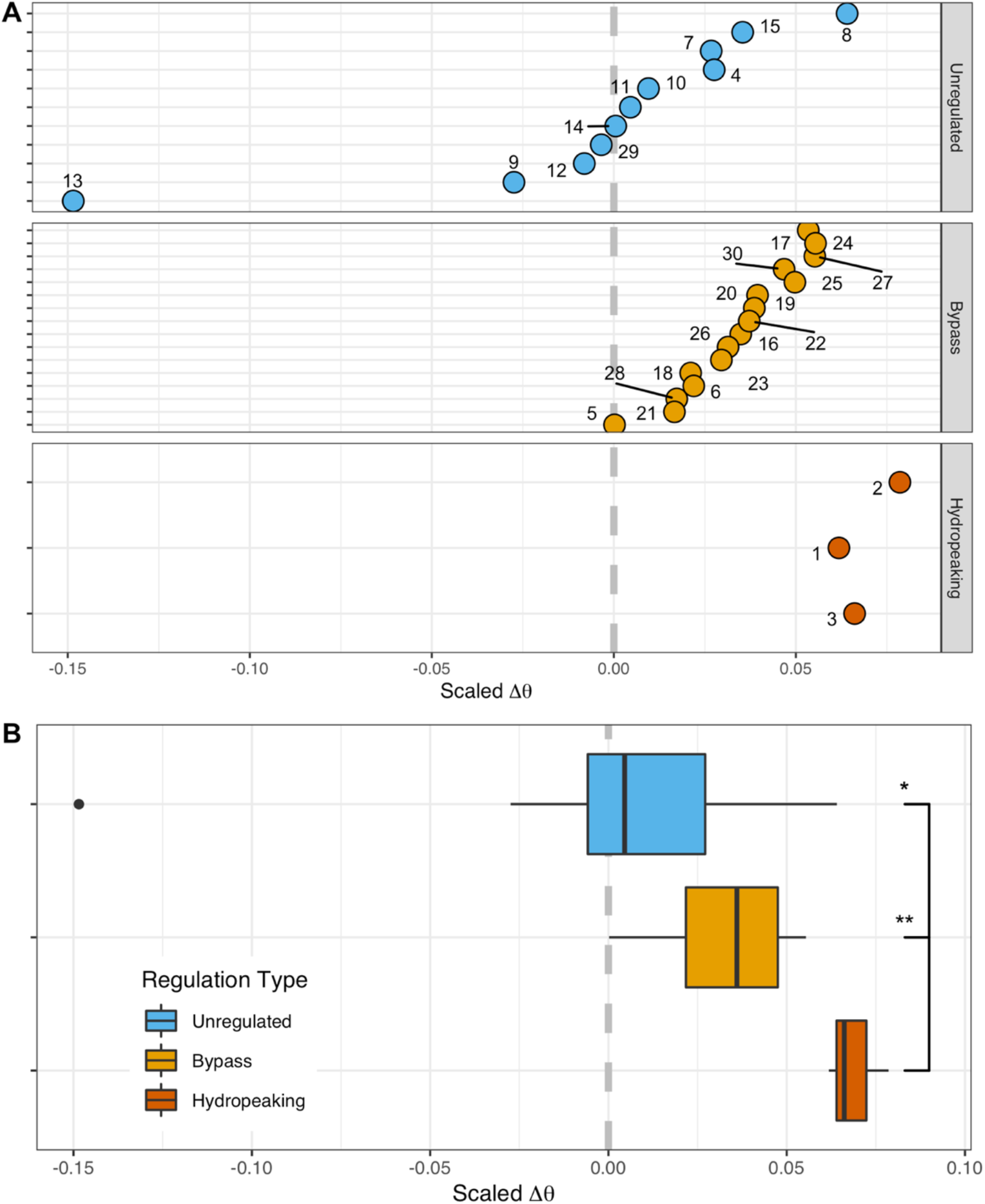
Relationship between river regulation and genetic diversity trajectory in *R. boylii*. A) Assessment of genetic diversity trajectories using scaled Δθ (θ_π_ − θ_S_ / [θ_π_ + θ_S_] / 2) for each sampling location; B) Boxplots of scaled Δθ and pairwise significance between regulation groups using a pairwise Wilcoxon rank sum test with Bonferroni correction (P < 0.05). Negative values represent trends of increasing genetic diversity, positive values represent trajectories of diversity loss, values near zero are stable.

## Discussion

Although massive parallel sequencing (MPS) technologies have the potential to facilitate collection of high-quality genetic data in virtually any species, a number of challenges still remain for many species including low quality or non-existent reference genomes, large/complex/repetitive genomes, and high cost of processing/sequencing in studies with many samples. Amphibians are particularly challenging as many species have very large genome sizes (Nunziata et al. 2017) for example, there are less than 10 frog reference genome assemblies available as of 2020 (Hellsten et al. 2010, Sun et al. 2015, Session et al. 2016, Hammond et al. 2017, Edwards et al. 2018, Rogers et al. 2018, Seidl et al. 2019) despite over 7,000 recorded frog species (AmphibiaWeb 2020). Our results demonstrate that Rapture (Ali et al. 2016) is a suitable method to rapidly and inexpensively discover a large number of loci in a frog species with a complex genome. In this study, we used new RAD sequencing and RAD capture (Rapture) methods (Ali et al. 2016) to generate high-quality genomic data suitable for discovering and genotyping many single nucleotide polymorphisms (SNPs) in *R. boylii*. Based on this dataset, we were able to successfully characterize patterns of genetic variation within *R. boylii* as well as design a set of RAD capture baits that can be used as a genetic monitoring resource for *R. boylii* (and likely other ranid species). This highlights that the collection of genetic information, even from large numbers of samples or in complex genomes, is no longer a limitation with current genomic methods such as RAD and Rapture.

Demographic connectivity is widely recognized as a fundamental driver of long-term population persistence (Fahrig and Merriam 1985, Taylor et al. 1993). Populations must adapt over time and connectivity is a major way to transfer genetic information. For example, previous studies have shown that adaptation can occur by spreading specific alleles across large geographic distances (Miller et al. 2012, Prince et al. 2017). In many regulated river reaches in the Sierra Nevada, *R. boylii* now occur in isolated locations, breeding in tributaries rather than mainstem habitats. However, since these frogs have the potential to move long distances (*R. boylii* have been observed moving over 1 km per day (Bourque 2008)), the extent to which current population connectivity has been lost due to river regulation was not understood. Examining pairwise F_ST_, revealed a major decrease in connectivity in populations in regulated systems, even with low intensity flow alteration (i.e., bypass reaches). The primary factor influencing genetic differentiation among these rivers is hydrologic alteration (Figure 3B). Thus, despite being able to move long distances, *R. boylii* have not been able to maintain population connectivity in rivers with altered flow regimes. This demonstrates that even in species that can move relatively long distances and pass potential physical barriers (e.g., infrastructure such as dams, canals, and reservoirs likely do not represent barriers to movement of *R. boylii*) loss of connectivity may still occur and can be revealed with genetic analysis.

Genetic diversity is also a critical component for long-term population persistence because it is closely related to the evolutionary capacity for adaptation to environmental changes (Lande and Shannon 1996, Frankham 2002, Hoffmann and Sgrò 2011, Ishiyama et al. 2015). In some cases, isolated populations can maintain genetic diversity when they are sufficiently sized (Whiteley et al. 2010), however, species of conservation concern typically have small and/or declining populations and thus may be susceptible to genetic diversity loss (Frankham 2002, Krohn et al. 2018). Throughout the Sierra Nevada, *R. boylii* have largely disappeared from regulated mainstem rivers, but the extent to which existing populations have been able to maintain genetic diversity is unclear. Strikingly, our analysis revealed that every single population within the regulated watersheds exhibits a trajectory of genetic diversity loss. Thus, genomic analysis of molecular variation provides a powerful lens to discover and assess trajectories of genetic diversity.

Our analyses, using metrics that serve as a reasonable proxy for genetic health, does not bode well for the long-term persistence of *R. boylii* populations in regulated rivers in the Sierra Nevada. Many of these *R. boylii* populations are already losing genetic diversity and the effects of inbreeding will likely exacerbate their problems given their small size and reduced connectivity. *Rana boylii* have evolved in river systems with consistent hydrologic seasonality and predictability, despite inter-annual variation. Flow regulation has altered patterns of hydrologic seasonality and predictability in many watersheds (Kupferberg et al. 2012). Long-term population persistence may still be possible if conservation efforts utilize methods that promote or maintain genetic health and increase population connectivity. For example, simulations by Botero et al. (2015) demonstrated adaptation persisted in modeled populations through large environmental changes—if phenotypic strategies were appropriate before and after the change—but modeled populations declined rapidly when the current strategy was a mismatch to the current environment. Thus, *R. boylii* conservation efforts should focus on river reaches where flow management may provide opportunities to more closely mimic local natural flow regimes and thus improve hydrologic connectivity.

Detecting evolutionary responses to within- and among-year changes in an ecological or hydrological context has previously been difficult. However, utilizing genetic data can fill these gaps and provide a highly informative process for identifying the impacts of anthropogenic and environmental change on the process of adaptation (Kahilainen et al. 2014, Botero et al. 2015). We demonstrate that an aquatic species that has adapted to local hydrology patterns shows poor genetic health (i.e., clear patterns of decreased connectivity and trajectories of genetic diversity loss). Our results highlight the impact of river regulation on aquatic organisms and their potential for long term persistence. In the future, similar genetic approaches could be used in many other contexts to explore the impacts of river regulation on aquatic organisms. Taken together, our results demonstrate that genetic monitoring can be a powerful tool for assessment of population health and should be a critical component of conservation management in aquatic organisms.

## Acknowledgements

This research builds on work from Amy Lind, Sarah Kupferberg, Jennifer Dever, and Sarah Yarnell. Many thanks to all of them for insight, and to many who helped collect/prepare samples: Corey Luna, Kristen Hein Strohm, Rick Wachs, and Isaac Chellman. This research could not have been conducted without access to specimens from Brad Shaffer at the Krebb Museum, and field samples from Sarah Mussulman, Caren Goldberg, and Mallory Bedwell. The authors have no conflicts of interest to declare.

## Data Archiving Statement

Should the manuscript be accepted, the data supporting the results will be archived in an appropriate public repository such as Dryad, and the data DOI will be appended to the end of the article.

